# MCL-1 as a molecular switch between myofibroblastic and pro-angiogenic features of breast cancer-associated fibroblasts

**DOI:** 10.1101/2025.02.10.637434

**Authors:** Chloé C. Lefebvre, Philippine Giowachini, Jennifer Derrien, Maxime Naour, Isabelle Corre, Elise Douillard, François Guillonneau, Mario Campone, Philippe P. Juin, Frédérique Souazé

**Affiliations:** Université de Nantes, INSERM, CNRS, CRCI2NA, F-44000 Nantes, France; Equipe labélisée LIGUE Contre le Cancer, Paris, France; SIRIC ILIAD, Nantes, Angers, France; Nantes Université, INRAE UMR1280, PhAN, IMAD, F-44000 Nantes, France; PROT’ICO - Plateforme Oncoprotéomique, Institut de Cancérologie de l’Ouest (ICO), Angers, France; ICO René Gauducheau, F-44800 Saint Herblain, France

## Abstract

Breast cancer-associated fibroblasts (bCAFs) comprise inflammatory CAFs (iCAFs), characterized by the secretion of pro-inflammatory cytokines, and myofibroblastic CAFs (myCAFs), distinguished by their high production of extracellular matrix and their immunosuppressive properties. We previously showed that targeting the anti-apoptotic protein MCL-1 in primary culture of bCAF derived directly from human samples reduces their myofibroblastic characteristics. We herein show by single-cell RNA-sequencing analysis of bCAFs that MCL-1 knock down induces a phenotypic shift from wound-myCAF to IL-iCAF, characterized by the upregulation of genes associated with inflammation as well as angiogenesis-related genes. *In vitro*, genetic and pharmacologic MCL-1 inhibition increases VEGF secretion by bCAFs, enhancing endothelial cell tubulogenesis. In a chicken chorioallantoic membrane (CAM) model *in ovo*, co-engraftment of breast cancer cells and bCAFs with reduced MCL-1 expression leads to heightened peritumoral vascular density, driven by VEGF. Mechanistically, the pro-angiogenic phenotype revealed by MCL-1 inhibition is dependent on BAX-BAK activity. It results in NF-κB activation, inhibition of which by a IKKβ inhibitor suppresses the transcription of VEGF and pro-inflammatory factors triggered by MCL-1 inhibition in bCAFs. Chemotherapy induces a downregulation of MCL-1 in bCAFs and it promotes a pro-angiogenic phenotype, counteracted by overexpressed MCL-1. Overall, these findings uncover a novel regulatory function of MCL-1 in determining bCAF subpopulation differentiation and highlight its role in modulating their pro-angiogenic properties, in response to treatment in particular.

## INTRODUCTION

Breast cancer-associated fibroblasts (bCAFs) are a predominant cellular component of the tumor stroma in breast cancers. Studies have identified a significant association between the abundance of bCAFs within tumors and unfavorable patient outcomes, underscoring their potential as therapeutic target ^1–3^. Single-cell transcriptomic analysis have identified two primary populations of bCAFs, namely inflammatory CAFs (iCAFs) and myofibroblastic CAFs (myCAFs), each comprising three distinct clusters. iCAFs are known for their extensive production of pro-inflammatory soluble factors like IL-6, CXCL12, CXCL1 and others that can attract immune cells on the tumor site and include subsets defined by specific functional pathways characterized by detoxification processes (Detox-iCAF), associated with interleukin signaling (IL-iCAF) and linked to IFNγ-mediated responses (IFNγ-iCAF). myCAFs are associated with a high contractile cytoskeleton linked to their pro-invasive features ^4,5^ and an important role in the production and organization of the extracellular matrix ^6,7^. In fact, the three myCAF clusters are characterized by a high expression of genes coding ECM proteins (ECM-myCAF), TGFβ signaling pathway (TGFβ-myCAF) or wound healing (Wound-myCAF) ^7^. Interestingly, Detox-iCAFs serve as precursors to both iCAFs and myCAFs. The Wound-myCAF state represents a transient and less differentiated intermediate between Detox-iCAF and ECM-myCAF, acting as a bridge to more specialized subtypes, including ECM-myCAF and IFNαβ-myCAF, as revealed by trajectory analyses ^8^. A spatial transcriptomic study showed that iCAFs are mainly found in the peritumoral stroma, while myCAFs are located in close contact with cancer cells ^8^.

Numerous studies have previously shown that CAFs promote tumor angiogenesis ^9–11^, cancer cell proliferation, survival and therapy resistance ^6,7,12,13^ in addition to metastases development ^14–16^. Treatment resistance can be mediated, in part, by the altered balance between pro- and anti-apoptotic expression of the members of the BCL-2 family (with BCL-2, BCL-xL and MCL-1 being the main anti-apoptotic proteins) ^17^. We previously demonstrated that bCAFs confer resistance to apoptosis in breast cancer cells through paracrine signaling by inducing overexpression of the anti-apoptotic protein MCL-1 ^18^. Targeting MCL-1 in breast cancers appeared to be a promising strategy, especially as we identified it as the most commonly expressed anti-apoptotic protein in bCAFs as well ^18^. We showed that targeting MCL-1 in bCAFs leads to non-lethal mitochondrial fragmentation accompanied by a loss of their myofibroblastic characteristics and pro-invasive abilities suggesting a role for MCL-1 in bCAFs plasticity ^19^.

MCL-1 is an essential regulator of mitochondrial homeostasis, mainly through its role in maintaining mitochondrial membrane integrity. Disturbances in this integrity influence various cellular processes, including metabolism, proliferation and apoptosis ^20^. Recent studies have also established a link between mitochondrial integrity and the regulation of inflammatory pathways ^21–23^. Which of these MCL1 functions contribute to the myofibroblastic features of bCAFs, and the underlying mechanisms, have remained elusive so far. In this study, we used genetic engineering of primary CAFs, single RNA sequencing and a whole set of *in vitro* and *in ovo* explorations. We show that MCL-1 maintains the myofibroblastic features of CAFs by preventing NF-KB driven acquisition of an inflammatory phenotype endowed with angiogenic properties. Moreover, chemotherapy triggers MCL-1 downregulation dependent angiogenic properties in bCAFs, defining MCL-1 as a chemotherapy actionable switch regulating stromal features.

## RESULTS

### Single cell RNA sequencing analysis reveals a transition from myofibroblastic bCAFs to an inflammatory phenotype associated with pro-angiogenic properties after MCL-1 gene silencing

For in depth study of the impact of MCL-1 targeting on the cellular composition of bCAFs, we performed a single cell transcriptomic analysis on three different patient derived primary cultures of bCAFs genetically engineered to be deficient in MCL-1 (bCAFsgMCL-1) or not (bCAFsgCTRL) using CRISPR/Cas9 technology (Figure 1A). After integration data and unsupervised clustering, we identified four distinct clusters, labeled 0 to 3 (Figure 1B). Enrichment analysis of gene sets preferentially expressed in each cluster indicated that phenotypic diversity in our primary cultures relied, at least in part, on differences in actors of Rho-GTPase cycle, metabolic process and VEGF-VEGFR signaling (cluster 0/2), cytoplasmic translation (cluster 1), mitotic progression (cluster 2) and protein phosphorylation (cluster 3) (Supplementary Figure 1A). Importantly, MCL-1 knock down enriched the representation in clusters 0 and 2 at the expense of clusters 1 and 3 (Figure 1C).

**Figure 1.**
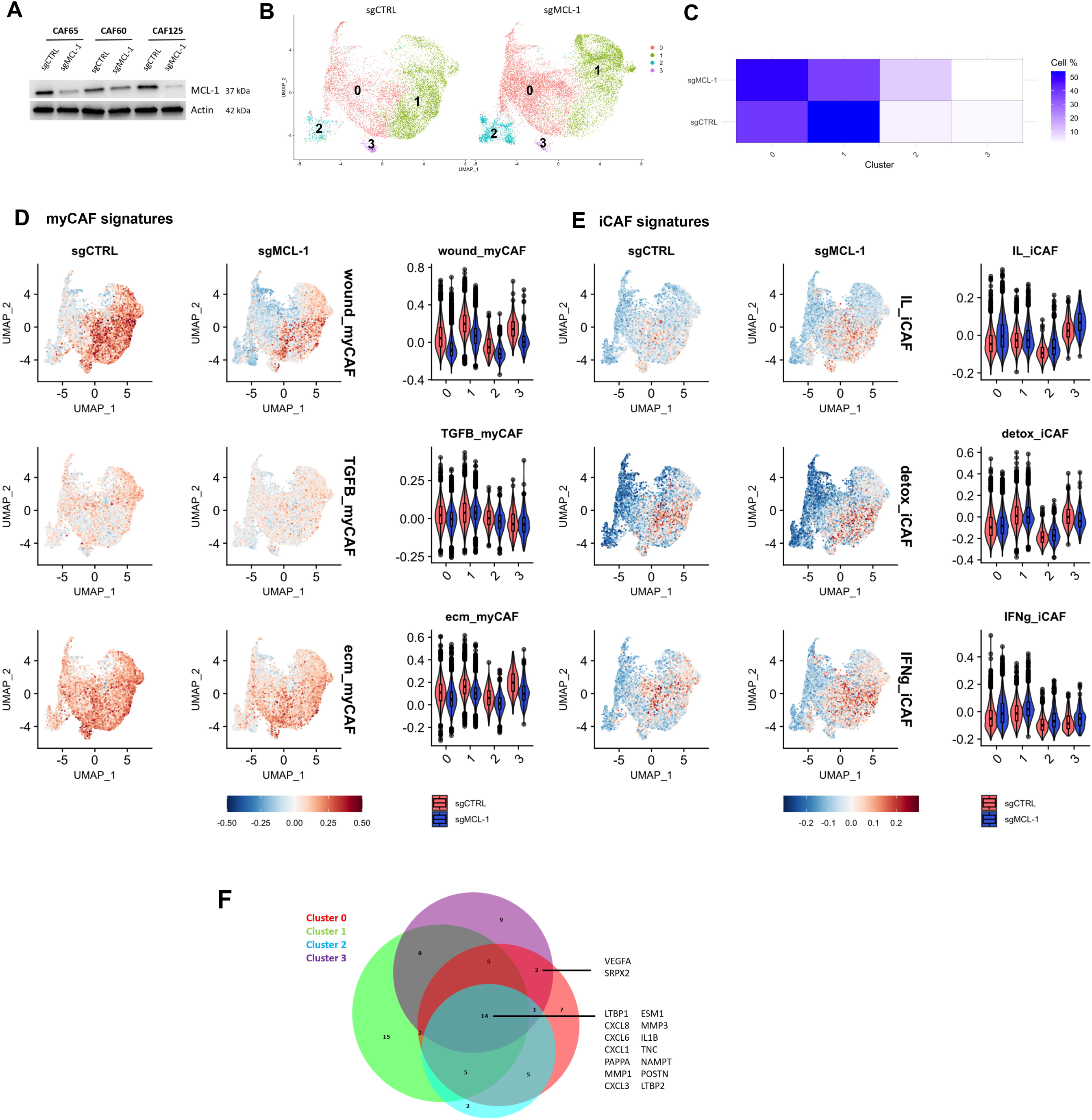
Single cell RNA sequencing analysis reveals a transition from myofibroblastic bCAFs to an inflammatory phenotype associated with pro-angiogenic properties after MCL-1 gene silencing. **A.** MCL-1 protein expression level in bCAFs after gene silencing evaluated using western blots. Actin expression was used as loading control (n=3). **B.** UMAP showing clusters of bCAF expressing MCL-1 (sgCTRL) or not (sgMCL-1)**. C.** Stack bar plot showing number of cells (left) and percent of cells (right) in each cluster under all conditions. **D. E.** (Top) Identity score of **(D)** myCAF signatures (wound-, TGFB- and ECM-myCAF) and **(E)** iCAF signatures (IL-, Detox- and IFNg-iCAF) of Kieffer et al ^7^ and (Bottom) violin plot showing signature score in each cluster in all conditions. **F.** Venn diagram of upregulated secreted factors (log2FC > 0.3, q-value < 0.01) in each cluster after MCL-1 gene silencing in bCAFs built with DeepVenn.

To further decipher how MCL-1 regulates myofibroblastic features ^19^, we evaluated six bCAF signatures defining myofibroblastic (wound-, TGFB-, ECM-phenotypes) myCAF and inflammatory (IL-, detox-, IFNg-phenotypes) iCAF as determined by Kieffer ^7^ in each single cell of our landscape. As expected from our culture method, our bCAF primary cultures expressed high levels of myCAF-associated genes. Myofibroblastic features were found across the four different clusters, with varying frequencies. In agreement with our preceding study, MCL-1 invalidation reduced myCAF scores. Yet the intensity of decrease differed between phenotypes and clusters. The wound myCAF was particularly sensitive to MCL-1 down-regulation while other myofibroblastic phenotypes somehow maintained in some clusters (TGFb in cluster 1 for instance) (Figure 1D). The high sensitivity of the wound phenotype is in agreement with its assigned plasticity ^8^. Strikingly, MCL-1 silencing also resulted in the acquisition of IL-iCAFs (otherwise rarely found in control bCAFs) in clusters O and 3 which showed the most dramatic drop in wound myCAF phenotype (Figure 1E). Taken together these results argue that loss of MCL-1 in bCAFs promotes a shift from myofibroblastic to inflammatory phenotype in plastic wound myCAFs.

To better characterize the inflammatory and secretory phenotype of bCAF following MCL-1 inhibition, we focused our differential analysis on the expression of genes encoding secreted factors in each cluster. Increased expression of genes encoding pro-inflammatory chemokines/cytokines such as CXCL8, CXCL6, CXCL1, IL1B after MCL-1 gene silencing was found in all clusters (Figure 1F). Strikingly a significant increase in VEGF-A upon MCL-1 silencing was detected in IL-iCAF-enriched groups 0 and 3 after MCL-1 inhibition hinting on a possible role for MCL-1 in preventing the pro-angiogenic properties of inflamed CAFs (Figure 1F and Supplementary Figure 1B).

### MCL-1 expression in CAFs is linked to their secretion of VEGF-A

To confirm a role for MCL-1 on the angiogenic features of bCAFs, we first examined both gene silencing (Figure 2A and B) and pharmacological inhibition of MCL-1 (Figure 2C and D) effects on expression and secretion by bCAFs of different angiogenesis-linked growth factors and cytokines (namely VEGF-A; FGF2, also known as FGF-basic; and ANGPT1). Either MCL-1 gene silencing or inhibition by S63845 ^24^ resulted in an upregulation of VEGF-A mRNA level (Figure 2A and C) and in an increase of its secretion by bCAFs (Figure 2B and D). Pharmacological inhibition of MCL-1 by S63845 also induced FGF2 mRNA level but had no effect on its secretion (data not shown). Notably, we found a significant negative correlation between MCL-1 expression in bCAFs and their VEGF-A secretion (Figure 2E), suggesting that differences in MCL-1 expression between patient derived primary bCAFs cultures might account, at least in part, for their differences in angiogenic properties. Targeting BCL-xL, another anti-apoptotic protein expressed by bCAFs ^18^, by either gene silencing (Supplementary figure 2A) or pharmacological inhibition with A1331852 had no detectable effect on pro-angiogenic factors mRNA levels or VEGF-A secretion (Supplementary Figure 2B, C, D, E). Similarly, we found no correlation between the level of BCL-xL expression in bCAFs and their ability to secrete VEGF-A (Supplementary Figure 2F). These results highlight a specific and unexpected role for the anti-apoptotic protein MCL-1 in the regulation of VEGF-A secretion by bCAFs.

**Figure 2.**
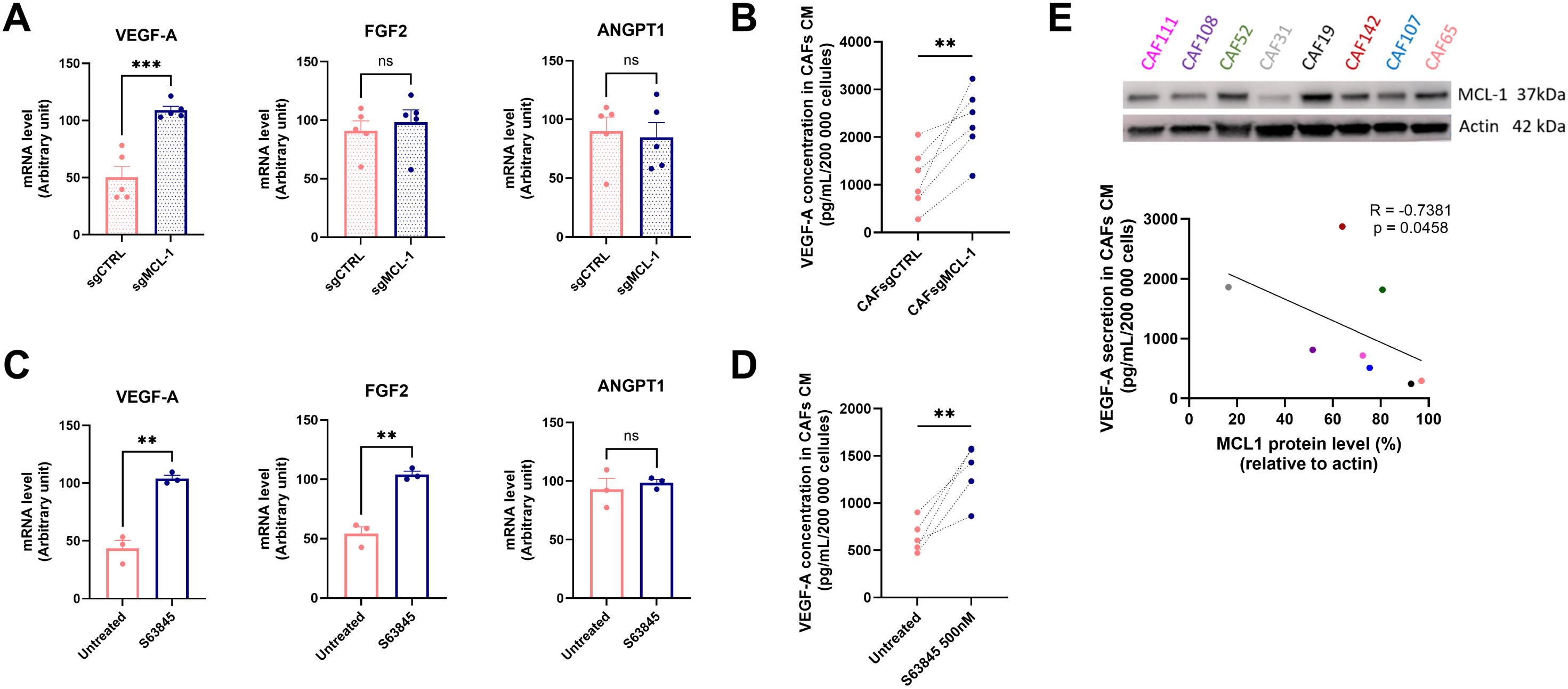
MCL-1 expression in CAFs is linked to their secretion of VEGF-A. **A.** qRT-PCR of VEGF-A, FGF2 and ANGPT1 mRNA in bCAFsgCTRL or sgMCL-1. Mean and SEM of five independent experiments are represented as relative quantity of mRNA. Student t-test, ***P < 0.001, ns: not significant. **B.** VEGF-A quantification by ELISA in conditioned media (CM) from bCAFs after MCL-1 gene silencing (bCAFsgMCL-1) or not (bCAFsgCTRL). The bCAFs CM were generated during 72h in EGM2 (Endothelial Cell Growth Medium-2) medium supplemented with 1% of FBS (n=6). Student t-test, **P < 0.01. **C.** qRT-PCR of VEGF-A, FGF2 and ANGPT1 mRNA in bCAFs treated or not by S63845 500 nM for 18 h. Mean and SEM of three independent experiments are represented as relative quantity of mRNA. Student t-test, **P < 0.01, ns: not significant. **D.** VEGF-A quantification by ELISA in CM of bCAFs treated or not by S63845 500 nM for 18 h Results were expressed as concentration (pg/ml) for 200 000 cells (n=5). Student t-test, **P < 0.01. **E.** Negative correlation (Spearman correlation coefficient r = − 0.7381; p value = 0.0458) between VEGF-A secretion level in bCAFs CM and MCL-1 protein level (relative to actin level) determined by western-blot analysis in 8 primary cultures of bCAFs.

### Targeting of MCL-1 in bCAFs promotes tubulogenesis of endothelial cells *in vitro* and angiogenesis *in ovo*

VEGF-A is a potent pro-angiogenic factor responsible for endothelial cell proliferation, survival and migration, which promotes angiogenesis during development, tissue vascularization and cancer progression ^25–27^. To evaluate the impact of MCL-1 targeting in bCAFs on angiogenesis, we generated conditioned media during 72h from bCAFs pre-treated with S63845 (CAF CM after S63845) or not (CAF CM Untreated) for 18h or from bCAFs silenced for MCL-1 (CAFsgMCL-1 CM) or not (CAFsgCTRL CM). We evaluated the capacities of endothelial cells to organize mini-vessels *in vitro* with tubulogenesis assay in response to bCAFs secretome (Figure 3A). Conditioned media from bCAFs pre-treated with S63845 significantly promoted tubulogenesis by inducing more master junctions, master segments and meshes in comparison with conditioned media from untreated bCAFs (Figure 3B). In parallel, conditioned media from bCAFs silenced for MCL-1 (CAFsgMCL-1 CM) also significantly enhanced the endothelial tubulogenesis in comparison with conditioned media from the control bCAFs (CAFsgCTRL) (Figure 3C).

**Figure 3.**
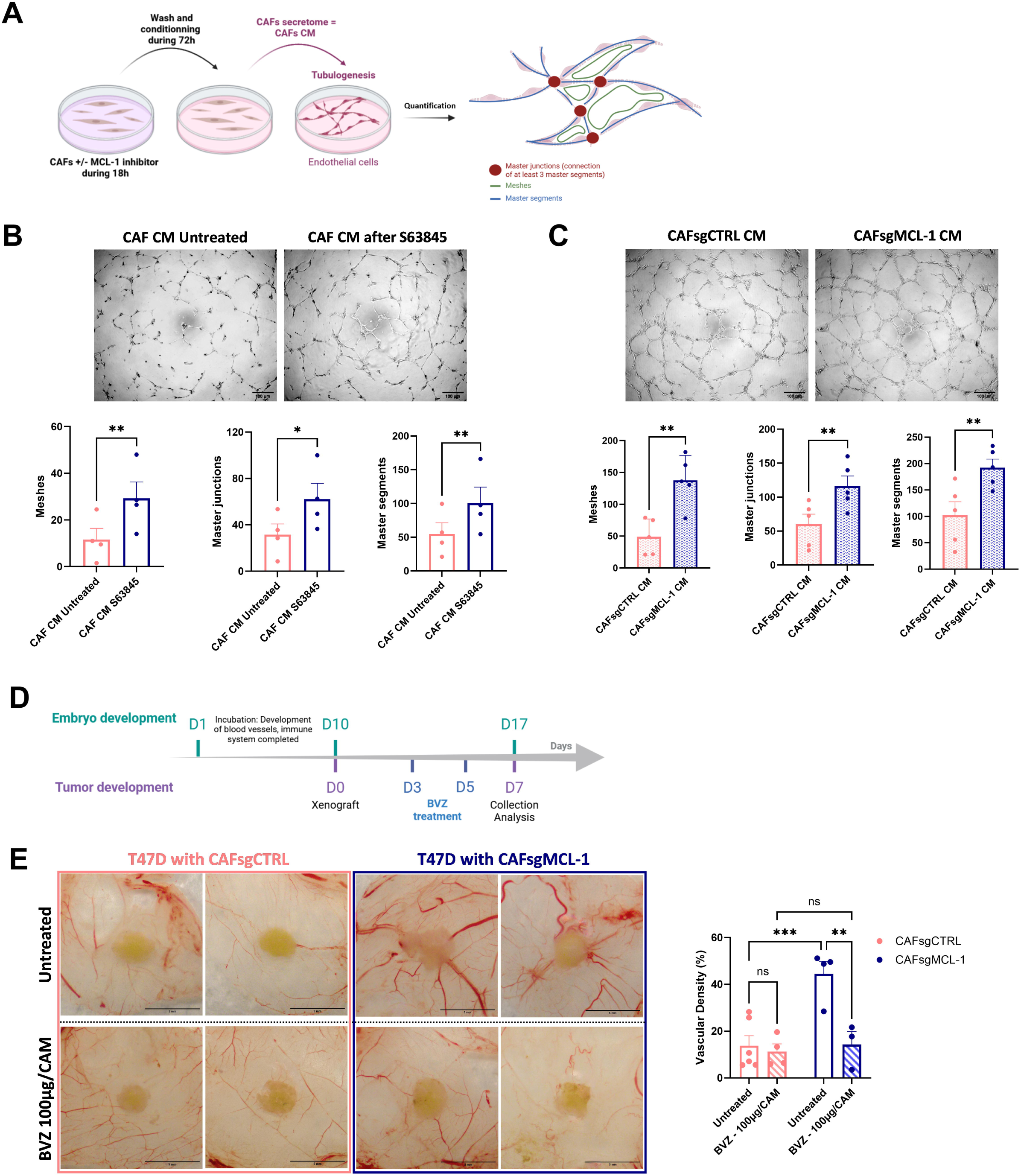
Targeting of MCL-1 in bCAFs promotes tubulogenesis of endothelial cells *in vitro* and angiogenesis *in ovo*. **A.** bCAFs were treated for 18 hours with MCL-1 inhibitor (S63845 500nM) or not. The treatment was removed and cells were rinsed and cultured for additional 72h in EGM2 (Endothelial Cell Growth Medium-2) medium supplemented with 1% of FBS. After 72h, the conditioned media (CM) were applied on HUVECs for tubulogenesis assay. **B.C.** (Top) Representative images of tubulogenesis assay of HUVECs under **(B)** CM from bCAFs treated or not with MCL-1 inhibitor (S63845 500nM) or **(C)** CM from bCAFs expressing MCL-1 or not. (Bottom) Quantification of meshes, master junctions and master segments of endothelial cells under **(B)** CM from bCAFs (untreated and S63845) n=4 or **(C)** CM from bCAFs sgCTRL or sgMCL-1 (n=5). Student t-test, *P < 0.5, **P < 0.01. **D.** Chronological timeline of CAM model experimentation. After 10 days of embryonic development (ED10), T47D luminal breast cancer cell line with bCAFs expressing or not MCL-1 (bCAFsgCTRL or bCAFsgMCL-1) were xenografted. The tumours were treated with VEGF inhibitor (bevacizumab, BVZ 100µg/CAM) every 2 days during one week until day 17 of embryonic development (ED17). **E.** (Left) Representative pictures of engrafted tumours on CAM at ED17 are shown (Top: untreated or Bottom: treated with bevacizumab, BVZ 100µg/CAM). (Right) Quantification of blood vessels density around the tumours (within a 5mm radius of the tumour). Two-way ANOVA, ***P < 0.001, **P < 0.01, ns: non-significant.

We investigated the impact of MCL-1 knock down in bCAFs on the angiogenesis *in ovo* using the chicken chorioallantoic membrane CAM. This vascularized alternative *in ovo* model allows the growth of numerous cell types ^28^. In this model, we performed xenografts of tumor models composed of T47D luminal breast cancer cells (chosen for their intrinsically low levels of VEGF-A secretion, approx. 50 pg/mL/200 000 cells, data not shown) mixed with primary bCAFs genetically modified for MCL-1 (CAFsgMCL-1) or not (CAFsgCTRL) (Figure 3D). Interestingly, one week after xenografts on CAM, we observed that the peritumoral vascular density is significantly increased around the tumours composed of bCAFsgMCL-1 compared to bCAFsgCTRL (Figure 3E, Top panel). Furthermore, these effects are annihilated with VEGF inhibitor treatment (BVZ) (Figure 3E, Bottom panel). These results highlight the strong involvement of MCL-1 in preventing VEGF-A dependant pro-angiogenic effect of bCAFs.

### VEGF-A secretion induced by MCL-1 inhibition in bCAFs involves BAX/BAK activity and is mediated by NF-kB

The canonical anti-apoptotic role of MCL-1 is due to its ability to sequester pro-apoptotic proteins, thereby inhibiting oligomerization of BAX-BAK pro-apoptotic effector proteins and preventing mitochondrial outer membrane permeabilization (MOMP). We assessed the secretion of VEGF-A triggered by S63845 in bCAFs lacking the pro-apoptotic proteins BAX-BAK as a result of a combined CRISPR approach (Figure 4A). As shown in figure 4B, S63845 induced no detectable VEGF-A secretion by bCAFs devoid of BAX-BAK. Inhibition of MCL-1 by S63845 did not trigger overt release of cytochrome-C from mitochondrial as assessed by FACS (Figure 4C) and the pan caspase inhibitor Q-VD-OPH did not mitigate S63845 mediated pro-angiogenic effect in bCAFs (Figure 4D). This suggest that VEGF-A secretion induced by MCL-1 inhibition in bCAFs involves non-lethal BAX and BAK activity.

**Figure 4.**
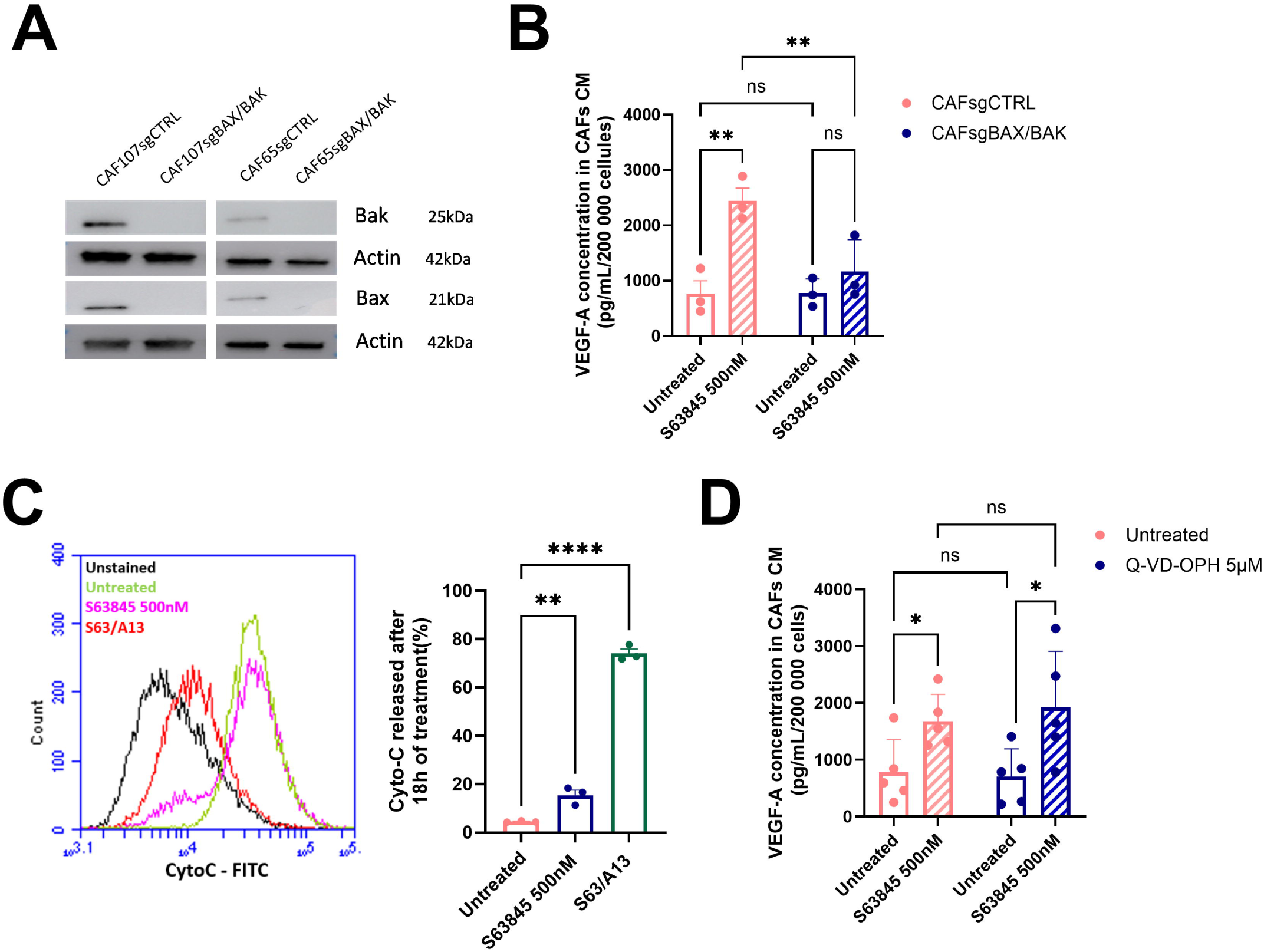
VEGF-A secretion induced by MCL-1 inhibition in bCAFs involves BAX/BAK activity and is mediated by NF-kB. **A.** BAX and BAK protein expression level in bCAFs after gene silencing evaluated using western blots. Actin expression was used as loading control (n=3). **B.** VEGF-A secretion analysed by ELISA in bCAFs CM expressing or not BAX/BAK (bCAFsgBAX/BAK) after MCL-1 inhibitor (S63845 500nM) or not (n=3). Two-way ANOVA, **P < 0.01; ns: non-significant. **C.** Cytochrome C released in bCAFs after 18h of treatment with S63845. Co-treatment with S63845 (500nM) and A1331852 (BCL-xL inh., 100nM) served as positive control of Cyto C release (n=3). Student t-test, **P < 0.01, ****P < 0.0001. **D.** VEGF-A secretion analysed by ELISA in bCAFs CM after MCL-1 inhibitor (S63845 500nM) in combination with pan-caspase inhibitor (Q-VD-OPH 5µM) (n=5). Two-way ANOVA, *P < 0.05; ns: non-significant.

To circumvent the phenotypic effect of targeting MCL-1 in bCAFs we performed a global proteomic analysis by mass spectrometry on four different primary cultures of bCAFs knock down for MCL-1 or not. We identified a total of 8184 proteins including 85 downregulated proteins and 196 upregulated proteins (log2FC > 0.58 and p-value < 0.05) after MCL-1 knock down in bCAFs (Figure 5A). Gene Ontology (GO) pathway enrichment analysis confirmed that MCL-1 gene silencing in bCAFs induced a decreased expression of proteins associated to their myofibroblastic phenotype and an increased expression of proteins implicated in inflammatory and innate immune response. These included TRADD, TRAF2, MIF, TNFAIP3 and CYLD related to the transcription factor NF-kB activity (Figure 5B). Moreover, TRANSFAC and JASPAR PWMs databases explorations indicated that a majority of upregulated proteins after MCL-1 gene silencing are transcriptional targets of RELA/NF-kB, which contributes to inflammatory phenotype acquisition (Figure 5C). This corroborates our single cell RNA sequencing analysis, as NF-kB-regulated genes are increased in clusters 0 and 3 after MCL-1 gene silencing (Figure 5D). MCL-1 targeting with S63845 induced a significant increase of NF-kB nuclear translocation in bCAFs (Figure 5E) and NF-kB activity blockade with an IKKβ inhibitor (AS602868) counteracted VEGF-A mRNA level increase after MCL-1 inhibition as well as the increase in CXCL-8, IL-1β and CXCL1 mRNA (Figure 5F). These results establish the implication of NF-kB signaling in MCL-1 targeting induced BAX/BAK dependent effects on VEGF-A synthesis (Figure 5G). Of note, we detected no effect of STING or TBK1 inhibition in bCAFs on the NF-kB dependent pro-inflammatory effects of MCL-1 targeting, ruling out the involvement of cGAS-STING/TBK1 dependent NF-kB activation downstream of BAX-BAK mitochondrial permeabilization ^29^ in this process (Supplementary figure 3A-E).

**Figure 5.**
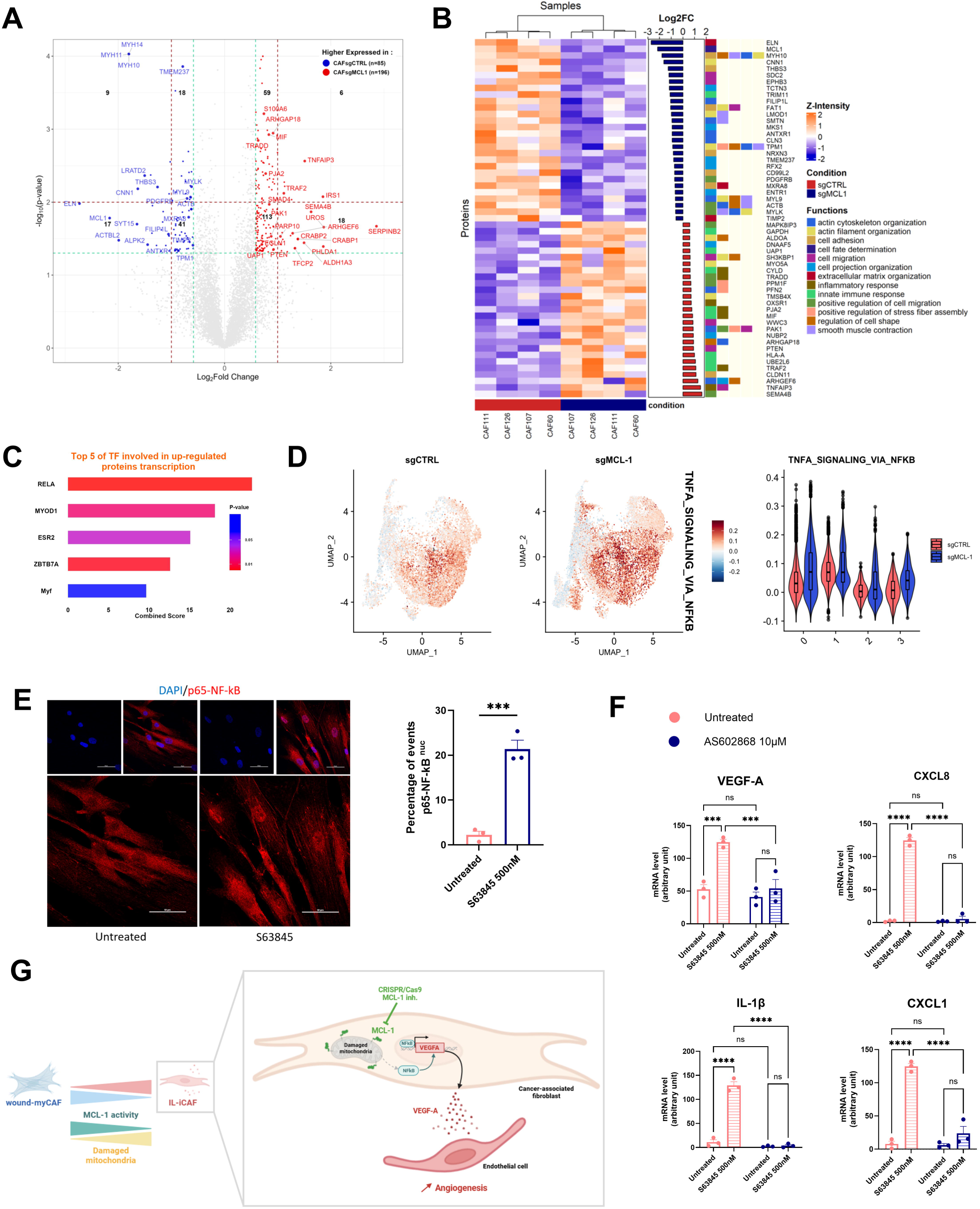
VEGF-A secretion induced by MCL-1 targeting is mediated by NF-kB in bCAFs. **A.** Volcano plot showing differential expressed proteins between bCAFs after MCL-1 gene silencing (bCAFsgMCL-1) or not (bCAFsgCTRL) (n = 4). The orange dots represent the upregulated expressed proteins; the blue dots represent the proteins whose expression is downregulated. **B.** A hierarchically clustered heatmap per sample showing differential expressed proteins associated to their biological process (Gene Ontology). Orange and blue represent up- and downregulated expression in bCAFs after MCL-1 gene silencing (sgMCL-1) or not (sgCTRL). Color density indicating proteins intensity levels. Log2FC of each protein was indicated on bar plot. **C.** Bar plot based on TRANSFAC and JASPAR PWMs databases (Enrichr) of the major transcription factors implicated in the transcription of the upregulated proteins after MCL-1 gene silencing in bCAFs. Combined score = -log(odds.ratio)xp-value. **D.** (Left) Gene set enrichment analysis with HALLMARK_TNFA_SIGNALING_VIA_NFKB on single cell RNA sequencing data of bCAFs expressing MCL-1 (sgCTRL) or not (sgMCL-1) and (right) violin plot showing signature score in each cluster under all conditions. **E.** Quantification (percentage of cells positive for nuclear p65) (right) and confocal image (left) of p65 staining (red) in bCAFs treated or not with S63845. The nuclei were counterstained with DAPI (blue). Scale bar = 50 μm. Student t-test, ***P < 0.001. **F.** qRT-PCR of VEGFA, CXCL8, IL-1β and CXCL1 mRNA in bCAFs after S63845 for 18h in combination or not with an IKKβ inhibitor (AS602868) (n=3). Two-way ANOVA, ****P < 0.0001, ***P < 0.001, ns: not significant. **G.** Graphical abstract showing bCAF phenotype dependence to MCL-1 activity. Targeting of MCL-1 induces mitochondrial dysfunction associated with nuclear translocation of NF-κB and VEGF-A transcription contributing to angiogenesis.

### Chemotherapy modulates the pro-angiogenic properties of bCAFs in an MCL-1-dependent manner

We finally analysed whether the function exerted by MCL-1 in bCAFs, described above, is modulated by chemotherapy. One standard of breast cancer treatment involves the combination of three chemotherapies: anthracycline (like doxorubicin), alkylating agents (like cisplatin) and anti-metabolites (like 5-fluorouracil). Such chemotherapeutic regimen did not induce cell death in bCAFs (Supplementary Figure 4A) but it drastically reduced MCL-1 protein levels while simultaneously increasing the pro-apoptotic NOXA protein in bCAFs (Figure 6A). Indeed, in many cell types, DNA-damaging chemotherapeutic agents stimulate transcription of NOXA ^30^ (Supplementary figure 4B) an endogenous inhibitor of MCL-1, that can interact with it to promote its degradation ^31–33^. BCL-XL level remained unchanged in all conditions (Figure 6A).

**Figure 6.**
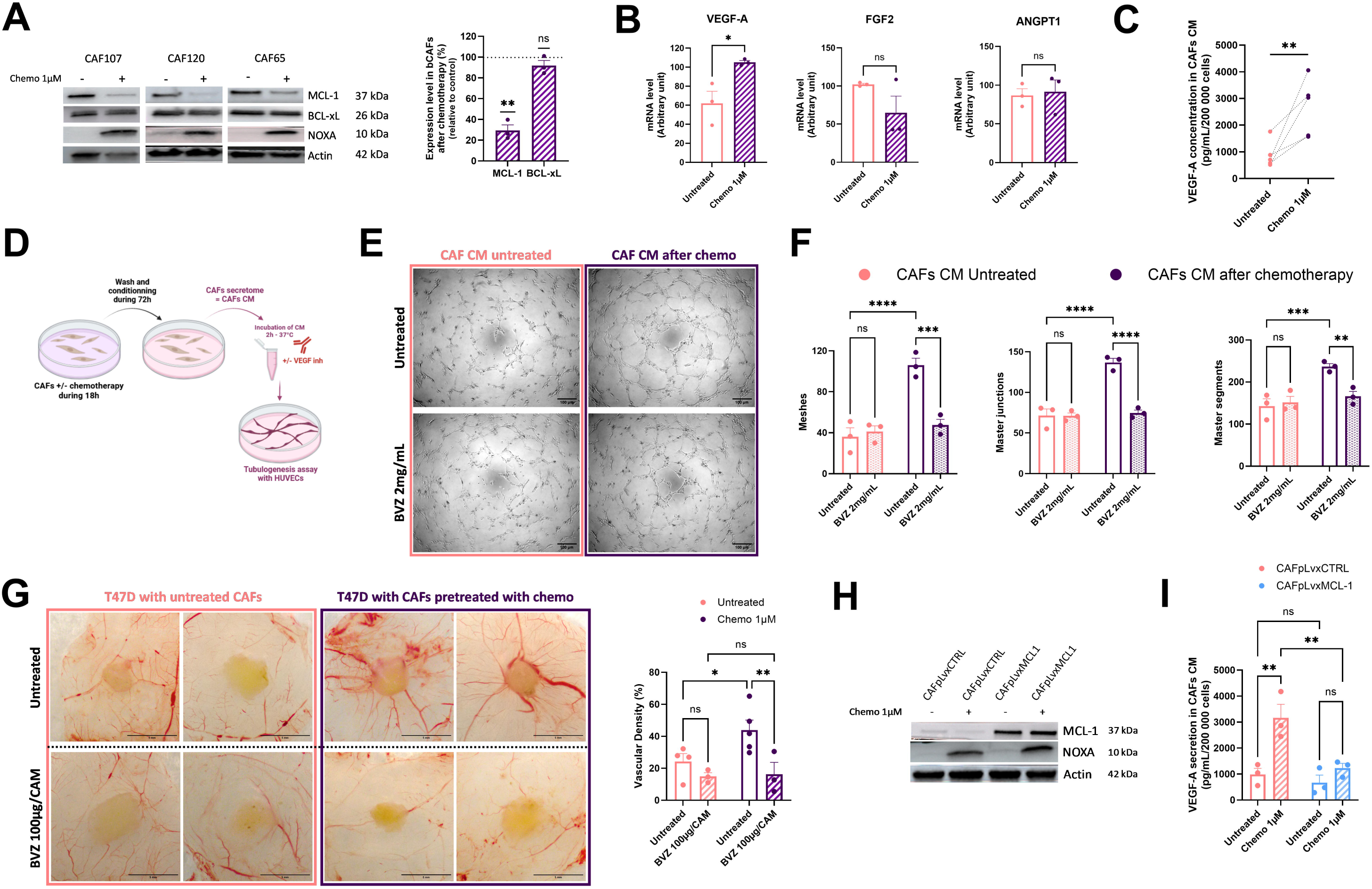
Chemotherapy modulates the pro-angiogenic properties of bCAFs in an MCL-1-dependent manner. **A**. (Left) Representative experiments of proteins expression level (MCL-1, BCL-xL and NOXA) in bCAFs after chemotherapy treatment (5-Fluorouracil 22µM, Cisplatin 11µM and Doxorubicin 1µM) or not evaluated using western blots, Actin expression was used as loading control. (Right) Quantification of the amounts of MCL-1 and BCL-xL protein band relative to Actin in bCAFs after chemotherapy treatment (n=3), results are expressed as a ratio relative to untreated bCAFs. One sample t-test, **P < 0.01, ns: non-significant. **B.** RT-qPCR of VEGF-A, FGF2 and ANGPT1 mRNA in bCAFs after chemotherapy treatment or not. Mean and SEM of three independent experiments are represented as relative quantity of mRNA. Student t-test, *P < 0.05, ns: not significant. **C.** VEGF-A was analysed by ELISA in bCAFs CM after chemotherapy. Student t-test, **P < 0.01. **D**. bCAFs were treated or not with chemotherapy for 18h. The treatment was removed and cells were rinsed and cultured for additional 72h in EGM2 medium supplemented with 1% of FBS. After 72h, conditioned media (CM) were recovered and a portion was incubated with VEGF inhibitor (BVZ, 2mg/mL) for 2 hours – 37°C before adding on HUVECs for tubulogenesis assay. **E.** Representative images of tubulogenesis assay of HUVECs cultured in conditioned media from bCAFs treated with chemotherapy or not +/- BVZ (2mg/mL). **F**. Quantification of meshes, master junctions and master segments of HUVECs cultured in conditioned media from bCAFs treated or not with chemotherapy +/- BVZ (2mg/mL) (n=3). Student t-test, **P < 0.01, ***P < 0.001, ****P < 0.0001, ns: non-significant. **G.** (Left) Representative pictures of engrafted tumours on CAM at ED17, tumours are composed of T47D cells and bCAFs pre-treated or not with chemotherapy for 18h before xenograft (Top). The tumours were treated with VEGF inhibitor (bevacizumab, BVZ 100µg/CAM) every 2 days during one week until ED17 (Bottom). (Right) Quantification of blood vessels density around the tumours (within a 5mm radius of the tumour). Two-way ANOVA, **P < 0.01, *P < 0.05, ns: non-significant. **H.** MCL-1 and NOXA proteins expression levels in bCAFs surexpressing MCL-1 (bCAFpLvxMCL-1) or not (bCAFpLvxCTRL) after chemotherapy were evaluated using western blots analysis. Actin expression was used as loading control. **I.** VEGF-A secretion was analysed by ELISA in bCAFs CM in the same conditions as (H) (n=3) Two-way ANOVA, **P < 0.01; ns: non-significant.

We found that MCL-1 protein level decrease by chemotherapeutic drugs was accompanied by an increase of VEGF-A mRNA without modification of FGF2 and ANGPT1 mRNA level in bCAFs (Figure 6B). VEGF-A secretion was increased by bCAFs after chemotherapy and accordingly, conditioned media from bCAFs in this condition promoted endothelial tubulogenesis in a BVZ sensitive manner (Figure 6C, D, E and F). Indeed, *in ovo*, we demonstrated that bCAFs pre-treated with chemotherapy before xenograft on CAM induced more blood vessels around the tumour compared to xenograft with bCAFs untreated (Figure 6G). Our results highlight the impact of chemotherapy on pro-angiogenic phenotype of bCAFs. Most importantly MCL-1 overexpression in bCAFs (Figure 6H and Supplementary figure 4C) counteracted the effect of chemotherapy on VEGF-A secretion (Figure 6I). Furthermore, chemotherapy-mediated pro-angiogenic effects of bCAFs were accompanied by an increase of CXCL8, IL-1β and CXCL1 mRNA level in bCAFs (Supplementary figure 4D). This argues that chemotherapy enables the pro-inflammatory and pro-angiogenic properties of bCAFs by boosting MCL-1 degradation.

## DISCUSSION

In this study, we highlight a phenotypic transition of bCAFs from a myofibroblastic to an inflammatory and pro-angiogenic phenotype and show that it is controlled by MCL-1 canonical activity. This emphasizes the role of MCL-1 in the overall organization of breast cancer ecosystems, in which it over-expression has frequently been associated with resistance to treatment ^18, 34^. This also puts forth a plasticity in bCAFs populations that could explain their phenotypic response to chemotherapy.

In breast cancer, two major pro-tumoral bCAFs subpopulations have been described including myCAFs associated with contractile and immunosuppressive phenotype and iCAFs known for their extensive production of inflammatory factors ^7^. We have previously shown that targeting the anti-apoptotic protein MCL-1 in bCAFs reduced their contractile and invasive capacities. Studies in cardiomyocytes reported similar findings, suggesting a role for MCL-1 in contractile cell function maintenance ^35^. Our single-cell RNA sequencing analysis of primary bCAF cultures following MCL-1 gene silencing demonstrates that the myofibroblastic wound-myCAF phenotype of bCAFs is highly dependent on MCL-1. Moreover, the loss of MCL-1 expression induces a phenotypic shift towards an inflammatory profile, particularly characterized by the acquisition of an IL-iCAF phenotype, incidentally underscoring the high plasticity of bulk bCAFs populations, even *in vitro*. In their recent study, Croizer et al ^8^ identified a differentiation pathway among bCAF subpopulations, with Detox-iCAF proposed as the origin of all other bCAF subtypes. In our work, wound-myCAFs were most affected by MCL-1 gene silencing, somehow consistent with their higher phenotypic plasticity compared to ECM-myCAFs and TGFβ-myCAFs that are at more advanced differentiation stages. Since MCL-1 knock down favors an IL-iCAF phenotype while exerting no detectable effect on a Detox-iCAF one, we propose that a novel MCL-1-dependent reversible differentiation pathway leading to the transition from IL-iCAFs to Wound-myCAFs exists. This is in agreement with Croizer’s study ^19^ showing a major role for YAP1 transcription factor activity in the maintenance of the myCAFs phenotype, as we showed that inhibition of MCL-1 in bCAFs leads to cytoplasmic retention of YAP1 ^19^. In prostate cancer, YAP1 has also been shown to act as a guardian of the myCAF phenotype by limiting the iCAF phenotype via inhibition of NF-κB activity ^36^. Acquisition of inflammatory/angiogenic features in plastic bCAFs upon MCL-1 invalidation might equally result from or cause the loss of (wound) myofibroblastic observed. However, MCL-1 appears to differentially regulate these two phenotypes at the molecular level. We showed that the loss of myofibroblast phenotype induced by MCL-1 inhibition (characterizing the first step in the transition from Wound-myCAFs to IL-iCAFs) was associated with mitochondrial fragmentation in a DRP1-dependent manner yet without the need for pore forming effectors of the mitochondrial outer membrane permeabilization (MOMP), BAX and BAK. In contrast, we herein showed that inhibition of MCL-1 leading to an IL-iCAF (secreting VEGF-A) phenotype required BAX/BAK activity. While this happens without overt cytochrome c release or caspase activation triggering cell death (^19^ and supplementary data), it nevertheless argues that MCL-1 prevents the acquisition of an inflammatory/angiogenic phenotype by bCAFs as a canonical guardian of their mitochondrial integrity. We thus propose that complete transition from Wound-myCAFs to IL-iCAFs upon MCL-1 invalidation comprise two molecular events at the mitochondrial level (mitochondrial fragmentation and permeabilization, respectively). Importantly, a direct effect of mitochondrial defects on the iCAF phenotype was observed in BRCA1-mutated cancers. In this context, germline BRCA1 mutations induce mitochondrial dysfunction, which amplifies the inflammatory phenotype of CAFs in triple-negative breast cancer, thereby facilitating angiogenesis and promoting tumor progression ^37,38^.

Our findings establish a connection between MCL-1 inhibition, NF-κB activation, and the acquisition of an inflammatory and pro-angiogenic phenotype by bCAFs. During MOMP, released mitochondrial contents like mtDNA can trigger inflammation via the cGAS-STING-NF-κB pathway ^29^. Our study demonstrated that S63845-induced transcription of VEGF-A, CXCL8, IL-1β and CXCL1 in bCAFs requires NF-κB activity but appears independent from STING/TBK1. Evidence suggests NF-κB activation can occur through NEMO (IKKγ) recruitment to ubiquitinylated mitochondria post-MOMP, forming a complex with IKKβ and IKKα, leading to IκBα degradation and NF-κB nuclear translocation ^22,23^. Further investigation is needed to clarify NF-κB activation mechanisms after S63845-induced mitochondrial damage in bCAFs. Interestingly, pro-inflammatory genes expressed by breast and ovarian CAFs are known to be NF-κB targets ^39^.

Our definition of MCL-1 as a key molecular switch between the opposing myofibroblastic and inflammatory phenotypes within bCAFs populations increase understanding of the effects of chemotherapy on bCAFs, which we show here (in the form of a doxorubicin, cisplatin, 5-fluorouracil treatment) to downregulate MCL-1 expression. The latter was shown to be down-regulated by anthracyclines, which are part of the chemotherapeutic regimen used in the treatment of breast cancer. We infer that chemotherapy-induced DNA damage and transcriptional arrest in bCAF leads to a decrease in MCL-1 levels, possibly further amplified by degradation due to increase NOXA levels. Importantly, we established that chemotherapy promotes VEGF-A secretion by bCAFs (which contributes to endothelial cell tubulogenesis *in vitro* and angiogenesis *in ovo)* unless MCL-1 is overexpressed. It is relevant to note here that prior reports have described MCL-1 as an inhibitor of chemotherapy-induced senescence across various models ^40,41^ and that chemotherapy induces in some cell types a senescence-associated secretory phenotype (SASP) which includes pro-angiogenic factors ^42,43^. Our study aligns to these observations by establishing a key role of MCL-1 is preventing the induction of an angiogenic phenotype in bCAFs upon chemotherapy ^40,41^.

MyCAFs (that rely on MCL-1 for maintenance) are described as immunosuppressive ^7^ and a decrease in their content post chemotherapy was shown to associate with T cell infiltration and to predict the efficiency of combined immunotherapy ^44^. In lung cancers in particular, low myCAF scores are associated with a better response to immunotherapies than stroma exhibit myCAF high score ^7^ Our data put forth that MCL-1 expression in bCAFs, prior and during chemotherapy, is one parameter whose clinical evaluation will document treatment response. The role of this protein in balancing the inflammatory and angiogenic versus myofibroblastic features of bCAFs nevertheless underlines possible adverse effects of its modulation by chemotherapy on the stroma. Recent studies have reported the pro-angiogenic power of iCAFs as well and their involvement in cancer progression ^38,45^. At clinical level, micro-vessel density is correlated with a poor prognosis with greater likelihood of metastatic disease and shorter survival rate for breast cancer patient ^46–48^. At pre-clinical and clinical levels, combination of immune checkpoint blockage and therapy based on targeting angiogenesis have beneficial synergically effects ^49^. Thus, the predictive value of MCL-1 stromal expression remains to be determined in specifically defined clinical contexts.

## METHODS AND MATERIALS

### Cell culture and reagents

Fresh human mammary samples were obtained from treatment naive patients with invasive carcinoma after surgical resection at the Institut de Cancérologie de l’Ouest, Nantes/Angers, France. As required by the French Committee for the Protection of Human Subjects, informed consent was obtained from enrolled patients and protocol was approved by Ministère de la Recherche (agreement no.: DC-2012-1598) and by local ethic committee (agreement no.: CB 2012/06). To isolate bCAFs from fresh samples, breast tissues were cut into small pieces in Dulbecco’s Modified Eagle Medium (DMEM Thermo Fisher Scientific) supplemented with 10% FBS, 2mM glutamine and 1% penicillin/streptomycin and placed in a plastic dish. bCAFs were isolated by their ability to adhere to plastic. After isolation, the fibroblasts were cultured in the same medium. Fibroblasts were used in the experiments before the ninth passage.

For the CRISPR Cas9-induced knock-out (KO) primary bCAFs, singleguide (sg) RNA sequences targeting human genes were designed using the CRISPR design tool (http://crispor.tefor.net). The following guide sequences were cloned in the lentiCRISPRV2 vector that was a gift from Feng Zhang (Addgene plasmid #52961): sgMCL-1: 5’-CTGGAGACCTTACGACGGGT-3’, sgBCL-xL: 5’-GCAGACAGCCCCGCGGTGAA-3’, sgBAK: 5’-GCCATGCTGGTAGACGTGTA-3’, sgBAX: 5’-AGTAGAAAAGGGCGACAACC-3’ and sgSTING: 5’-GCAGGCACTCAGCAGAACCA-3’. MCL-1 overexpressing bCAFs were established by viral infection with retroviruses containing vector coding for MCL-1 (pLVX-EF1alpha-IRES-Puro). Empty vector was used as control. Cells were selected using 1 μg/ml puromycin and protein extinction were confirmed by immunoblot analysis.

The human breast cancer cell lines T47D were purchased from American Type Culture Collection (Bethesda, MD, USA) and was cultured in RPMI medium supplemented with 10% FBS and 2mM glutamine.

Human umbilical vein endothelial cells (HUVECs) were freshly isolated from umbilical cords obtained from Nantes Hospital Maternity from different donors with informed consent from all subjects, as previously described ^50^ (De Bock, 2013). Endothelial cells (ECs) were routinely cultured in Endothelial basal medium (EGM2; containing 2% fetal bovine serum (FBS); PromoCell, Heidelberg, Germany) supplemented with Endothelial growth medium supplement pack (ECGM-2; PromoCell, Heidelberg, Germany) and 100 IU/mL penicillin and 100 mg/mL streptomycin on 0,1 % gelatin-coated flasks. Cells were used from passage 1 to 6. All cells were cultured at 37 []C and 5% CO_2_.

### Treatments

Doxorubicin (Selleckhem #S1208), Cisplatin (Selleckhem #S1166) and 5-Fluorouracil (Selleckhem #S1209) were used alone or mixed at 1μM, 11μM and 22μM respectively to obtain chemotherapy treatment. A1331852 (MedChemExpress #HY-19741), S63845 (ChemieTek #CT-S63845), Q-VD-OPH (R&D Systems, Abindgon, UK), VEGF inhibitor (Bevacizumab, Aybintio, Samsung Bioepis), ProtacTBK1 (Biotechne, 7259), ProtacCTRL (TBK1 Control Protac, Biotechne, 7260) and AS602868 (Merck-Serono International SA) were used at indicated concentrations.

bCAFs are still treated for 18h in DMEM containing 1% FBS. For TBK1 degradation experiments, S63845 was added for 18h after a 3h incubation with Protac. For experiments with pan-caspase inhibitor, QVDOPH was used for 18h in combination or not with S63845 or chemotherapy. QVDOPH was also added during the 72h of conditioning in EGM2.

### Single-cell RNA sequencing

Single cell RNA sequencing was performed using the Chromium Single-Cell 3 ‘v3.1 kit from 10X Genomics, following the manufacturer’s protocol. The libraries were sequenced on the NovaSeq 6000 platform (Paired-end, with 28 bp for Read1 and 90 bp for Read2). The raw BCL files were demultiplexing and aligned to the reference genome (refdata-cellranger-GRCh38-3.0.0) using the Cell Ranger Software Suite (v.7.0.1).

### Preprocessing and Quality Control of Single-Cell RNA Sequencing Data

The raw data were extracted from 10X format files using the *Read10X* function from the Seurat package (v4.4.0) in R v.4.1.1. Quality control filtering was applied to the feature-barcode gene expression matrix to retain only high-quality cells. Cells were selected based on the following criteria: more than 5,000 unique molecular identifiers (UMIs), more than 2,000 detected genes, and less than 20% of reads mapped to mitochondrial genes to exclude potentially dead or dying cells. Doublets were detected using the scDblFinder R package (v.1.6.0). A score was computed for each cell, and doublets were identified by applying the threshold defined by the scDblFinder package. Detailed quality control metrics are provided in Supplementary Table 1.

### Single-cell RNA sequencing analysis

Data normalization for each dataset was performed using the *NormalizeData* function from the Seurat package. Variable features were identified using the vst method, selecting the 2,000 most highly variable genes per dataset. To integrate the datasets, features that were repeatedly variable across all datasets were selected using the *SelectIntegrationFeatures* function. Data were then scaled using the ScaleData function, and cell cycle effects were minimized by regressing out S and G2M phase scores. To perform dataset integration, anchors were identified using the *FindIntegrationAnchors* function, followed by integration of the data with the *IntegrateData* function, creating an integrated assay.

Principal Component Analysis (PCA) was performed using the *RunPCA* function, and Louvain graph-based clustering was applied to the first 30 principal components. The resulting data were visualized using Uniform Manifold Approximation and Projection (UMAP) with the *RunUMAP* function. Distinct clusters were identified using the *FindNeighbors* and *FindClusters* functions, with a resolution of 0.1.

For each defined cluster, gene marker analysis was conducted using the *FindAllMarkers* function with the following parameters: only.pos = TRUE, min.pct = TRUE, and logfc.threshold = 0.25. Finally, signature scores were calculated using the *AddModuleScore*function.

For pathway enrichment analysis, we used fgsea package (v1.33.1), marker genes were identified using the FindAllMarkers function with parameters min.pct = 0 and logfc.threshold = 0. Genes were then ranked according to their differences in gene expression to the other clusters. Enrichment was assessed using a set of pathways that were selected by analysis of genes preferentially expressed in each cluster.

The list of genes encoding secreted factors was based on the UniProtKB list of proteins located outside cell membranes (Cellular component - Secreted). A manual curation process, based on literature, was performed to remove certain genes not coding for extracellular proteins.

### Proteomic analysis by LC-MS/MS

The pelleted cells were quickly thawed and proteins were concomitantly extracted, denatured and thiol groups were chemically reduced in 200µL of 0.1% Rapigest-SF® (Waters), 5mM DTT and 50mM ammonium bicarbonate pH 8.5, in a thermomix set at 95°C for 30min under 750 rpm shaking. Sonication was performed twice 30 seconds using an ultrasonic processor probe sonicator (130W, 20 KHz) at 20% power (Thermo Fisher Scientific). Alkylation of free thiols was performed by adding S-Methyl methanethiosulfonate (10 mM final concentration from Sigma) with a further ten-minute incubation at 37°C and 750 rpm shaking. 5µg porcine trypsin (Sciex) was used to generate bottom-up LC MS-compatible peptides, and incubated at 37°C overnight. Resulting peptides were then cleared from potential insoluble material by centrifugation at 16000g for 15 minutes. The supernatant-containing peptides was desalted using in-house pipette tips-packed with C18 described as “Stage-tips” procedure ^51^. Peptides were then eluted and dried in a vacuum centrifuge concentrator (Savant SPD121P from Thermo), resuspended in 42µL of 0.1% Formic Acid (FA) and submitted to microBCA peptide assay (Thermo) to allow the injection of 200ng of peptides to be injected in LC MSMS in DIA-PASEF mode as follows: 2µL of each sample was loaded and separated on a C18 reverse phase column (Aurora series 1.6µm particles size, 75µm inner diameter and 25cm length from IonOptics, Fitzroy Australia) using a NanoElute LC system (Bruker, Bremen Germany). Eluates were electrosprayed into a timsTOF Pro 2 mass spectrometer (Bruker) for the 60 min duration of the hydrophobicity gradient ranging from 99% of solvent A containing 0.1% FA in milliQ-grade H2O to 27% of solvent B containing 80% ACN plus 0.1% FA in mQ-H2O. The mass spectrometer acquired data throughout the elution process and operated in data-independent analysis (DIA) with PASEF-enabled method using the TIMS-Control software (Bruker). Samples were injected in batch replicate order to circumvent possible interferences of technical biases.

The m/z acquisition range was 250–1201 with an ion mobility range of 0.7–1.25 1/K0 [V s/cm2], which corresponded to an estimated cycle time of one second. DIA-PASEF windows, and collision energy were also left to default with a base of 0.85 1/K0 [V s/cm2] set at 20 eV and 1.30 1/K0 [V s/cm2] set at 59 eV. TOF mass calibration and TIMS ion mobility 1/K0 were performed linearly using three reference ions at 622, 922, and 1222 m/z (Agilent Technology, Santa Clara, CA, USA).

### LC-MS/MS raw data analysis

The raw data were extracted, normalized and analyzed using Spectronaut® 18.0.230605.50606 (Biognosys) in DirectDIA+ mode, which modelized elution behavior, mobility and MS/MS events based on the Uniprot/Swissprot sequence 2020 database of 20,365 human proteins and 381 entries of common contaminant proteins. Protein identification false discovery rate (FDR) was restricted to 1% maximum, with a match between runs option enabled, and inter-injection data normalization. The enzyme’s specificity was trypsin’s. The precursor and fragment mass tolerances were set to 15 ppm. Acetyl (Protein N-term) and oxidation of methionines were set as variable modifications while thiol groups from cysteines were considered completely alkylated by methylation. A minimum of two ratios of peptides was required for relative quantification between groups. Protein quantification analysis was performed using Label-Free Quantification (LFQ) intensities. Spectronaut statistical tools were used to visualize data, assess quality and quantitatively compare datasets. The resulting proteins LFQ values were log2(1.5) transformed and importantly a minimum of 5% of available p-values in at least one group was needed to include proteins in the differential abundance analysis. The proteomic analysis was conducted using a custom-developed R script (v.4.4.1). Gene Ontology (GO) terms related to “Biological Process” were assigned using the Ensembl database via the biomaRt R package (v.1.0.7) (https://cran.r-project.org/web/packages/biomartr/index.html). Duplicate proteins were then filtered by retaining only the gene with the lowest p-value and we considered that a protein was expressed when it was present in at least 3 out of 4 samples in each condition. A manual curation process, informed by a thorough review of relevant literature, was performed to assign some proteins to relevant Biological Process categories. Volcano plot was generated using the ggplot2 package (v.3.5.1), while heatmap was built with ComplexHeatmap package (v.2.20.0) (https://bioconductor.org/packages/release/bioc/html/ComplexHeatmap.html) using euclidean distance and ward D2 clustering method.

### RNA isolation and quantitative real-time PCR

Total RNA was isolated using Nucleospin RNA (Macherey-nagel, Hoerdt, France) and transcribed into cDNA by Maxima First Strand cDNA synthesis Kit (Thermo scientific). Quantitative RT-PCR (qPCR) was performed using the EurobioGreen qPCR Mix Lo-Rox with qTOWER (Analityk-jena, jena, Germany). Reaction was done in 10 μl final with 4 ng RNA equivalent of cDNA and 150 nM primers. Relative quantity of mRNA was estimated by Pfaffl method ^52^ and normalized on the average relative quantity of one housekeeping gene.

RPLP0 5’-AACCCAGCTCTGGAGAAACT/CCCCTGGAGATTTTAGTGGT-3’

GAPDH 5’-CAAAAGGGTCATCATCTCTGC/AGTTGTCATGGATGACCTTGG-3’

FGF2 5’-CTTCCTGCGCATCCACCCCG/AGCCAGGTAACGGTTAGCACACA-3’

ANGPT1 5’-GCTCCACACGTGGAACCGGA/CCAGCATGGTAGCCGTGTGGT-3’

VEGF-A 5’-AAGGAGGAGGGCAGAATCAT/CCAGGGTCTCGATTGGATGG-3’

IL-1β 5’-TGGCAATGAGGATGACTTGT/GGAAAGAAGGTGCTCAGGTC-3’

CXCL8 5’-AAGCTGGCCGTGGCTCTCTTG/TTCTGTGTTGGCGCAGTGTGGT-3’

CXCL1 5’-AACAGCCACCAGTGAGCTTC/GAAAGCTTGCCTCAATCCTG-3’

### Immunoblot analysis

Cells were re-suspended in lysis buffer (1% SDS; 10 mM EDTA; 50 mM TrisHcl pH 8.1; 1 mM PMSF; 10 μg/ml aprotinin; 10 μg/ml leupeptin; 10 μg/ml pepstatin; 1 mM Na3VO4 and 50 mM NaF). For western blotting, following SDS–PAGE, proteins were transferred to 0.45 µM nitrocellulose membranes using Trans-Blot® Turbo™ Transfer System Cell system (Bio-Rad). The membrane was then blocked in 5% nonfat dry milk TBS 0.05% Tween 20 and incubated with primary antibody overnight at 4 °C. Blots were incubated with the appropriate secondary antibodies for 1 h at room temperature and visualized using the Fusion FX (Vilber). The used primary antibodies were anti-MCL-1 (Cell Signaling, 94296, 1/1000), anti-Bax (Cell Signaling, 5023, 1/1000), anti-Bak (Cell signaling, 1210, 1/1000), anti-BCL-xL (Cell Signaling, 2764, 1/1000), anti-NOXA (Abcam, ab13654, 1/500), NFkB-p65 (Cell Signaling, 8242, 1/1000), p-NFkB-p65 (Cell Signaling, 3033, 1/500), TBK1 (Cell Signaling, 3504, 1/1000), STING (Cell Signaling, 13647, 1/1000), pH2AX (Millipore, 05-636, 1/1000) and anti-β-actin (Millipore, MAB1501R, 1/2000).

### ELISA

For the preparation of conditioned media (CM), 170 000 bCAFs were seeded in one well of a 6-well plate and were treated as indicated during 18h in DMEM 1% FBS. The cells are washed with PBS and cultured in EGM2 supplemented with 1% FBS for additional 72 hours. CM were then collected and centrifuged (2000 rpm, 10 min). Levels of VEGF-A (446504, Biolegend) in cell supernatants were determined according to the manufacturer’s protocol.

### Tubulogenesis assay

For tubulogenesis assay, wells of a 96-well plate were coated with 40µL of Matrigel (Cultrex Reduced Growth Factor Basement Membrane Extract, Type R1, # 3433-010-R1, Bio-techne) and allowed to polymerize for 30 min at 37°C. 15 000 HUVECs were seeded per well with bCAFs conditioned media (generated with the same protocol as for ELISA) for 6 hours. For experiments with VEGF inhibitor (Bevacizumab, Avastin), bCAFs CM were pre-incubated with antibody at 2mg/mL for 2h at 37°C before the tubulogenesis assay. The images were acquired on an EVOS XLCore microscope with a 4x objective. For tube quantifications, images were processed in a blind manner using ImageJ software (NIH). All conditions are performed in duplicate and the whole wells area was used for quantification. A representative area of interest of well was shown in picture in figures.

### Chick embryo Chorioallantoic membrane (CAM) model

Fertilized chicken eggs were purchased from EARL LES BRUYERES (28190 DANGERS, France), and incubated for 9 days at 37[]C at 55% relative humidity. At day 9 of their embryonic development (ED9), a window of an ∼1 cm-diameter was drilled on top of the air chamber of the eggshell. At ED10, 1x10^6^ T47D cell line and 1x10^6^bCAFs (ratio 1:1) were suspended in 25 μL DMEM medium containing 1% fetal bovine serum, 100 U/mL penicillin and streptomycin, and 25 μL Matrigel (Cultrex Reduced Growth Factor Basement Membrane Extract, Type R1, # 3433-010-R1, Bio-techne). The mix was incubated for 15 minutes at 37[]C and subsequently implanted into the CAM of each egg. The tumours were treated with a VEGF neutralizing antibody (Bevacizumab, 100µg/CAM) at 3 and 5 days after inoculation. Seven days after implantation (ED17), chick embryos were killed by decapitation. For experiments with chemotherapy, bCAFs were pre-treated during 18h with chemotherapy in DMEM supplemented with 1% of FBS before their xenograft in CAM. At the end of experiments, tumours were excised, photographed with a Canon EOS 250D camera and carefully weighed to determine their mass. The vascular density within a 5mm radius around the tumours was quantified on Fiji software using the formula: vessel area/total area * 100%.

### Cytochrome C release

bCAFs were fixed and permeabilized using FIX & PERM Cell Fixation and Permeabilization Kits (00-5523-00, ebioscience, San Jose, CA, USA) for 15min – RT. After PBS wash and centrifugation, cell suspension was incubated with AlexaFluor®488-conjugated human Cyt-C antibody (BD Pharmingen, 560263, 1/50) for 45min - RT. Alexa Fluor® 488 Mouse IgG1, κ Isotype Ctrl (Fc) Antibody (Biolegend, 400129) was used at the same concentration. Fluorescence intensity was measured on BD Accuri™ C6 (BD Biosciences).

### Immunocytochemistry

Cells were fixed in PBS containing 4% paraformaldehyde/4% sucrose for 15 min. Cells were permeabilized for 7 min at RT in 0.25% Triton-X-100 in PBS, washed twice with PBS, and blocked for 45 min at RT in PBS containing 10% BSA. Cells were incubated overnight at 4°C with NFkB-p65 (Cell Signaling, 8242, 1/200) diluted in PBS containing 3% BSA. After washing, cells were incubated for 45min at 37 °C with the appropriate Alexa 546-conjugated secondary antibody diluted in PBS containing 3% BSA. Cells were washed with PBS, incubated with DAPI diluted in PBS (Invitrogen, 1/1000) for 10min and mounted with ProLong Diamond Antifade Reagent (Invitrogen, Carlsbad, CA, USA). Fluorescence images were acquired with Nikon A1 Rsi Inverted Confocal Microscope (Nikon, Tokyo, Japan) with NIS-Elements software (Nikon). Images set position were generated randomly. Around fifty to one hundred fifty cells were analysed per condition. Counting events were done manually through NIS-Elements software (Nikon).

### Statistical analysis

Student’s t-test was used for statistical analysis in experiments with two groups, one sample t-test for statistical analysis in experiments with one group and Two-way analysis of variance (ANOVA) was used for statistical analysis for overall condition effects with GraphPad Prism 10.2 Software. Spearman’s correlation test was used for correlations. All data are presented as mean ± SEM of at least three independent experiments. The symbols correspond to a P-value inferior to *0.05, **0.01, ***0.001, and **** 0.0001.

## DATA AVAILABILITY

The data that support the findings of this study are available from the corresponding author, [FS], upon reasonable request. The mass spectrometry proteomics data are available via ProteomeXchange with identifier PXD058906.

## ACKNOWLEDGEMENTS

We would like to thank members of the “Stress adaptation and tumour escape” laboratory for their support, in particular Fétiveau A. for the design and construction of the molecular plasmid vector and Duarte L. for her help with experimental manipulations on the CAM experiments. We thank Drs Tartare-Deckert S. and Derangeon M. for their fruitful discussions and suggestions. Our sincere thanks go to Drs. Thirouard L. and Chiron D. for their technical advice on the CAM experiments. We would like to thank Treps L., Kreijbich M. and Cotinat M. for their technical support and discussions, and the members of the Gavard J. and Bidère.N team for supplying reagents and technical advice. We are most grateful to Boissard A. and Henry C. from the PROT’ICO core facility, Institut de Cancérologie de l’Ouest (Angers, France). We acknowledge the MicroPICell core facility (SFR Bonamy, BioCore, Inserm UMS 016, CNRS UAR 3556, Nantes, France), member of the Scientific Interest Group (GIS) Biogenouest, IBISA, and the national infrastructure France-Bioimaging supported by the French national research agency (ANR-10-INBS-04). We thank Genomics Core Facility GenoA, member of Biogenouest and France Genomique and to the Bioinformatics Core Facility BiRD, member of Biogenouest and Institut Français de Bioinformatique (IFB) (ANR-11-INBS-0013 for the use of their resources and their technical support.

## AUTHOR CONTRIBUTIONS

C.C.L., P.G., I.C., E.D. and F.S. conducted experiments. C.C.L., I.C., P.P.J and F.S. designed the experiments. C.C.L., P.G., J.D., M. N., I.C., F. G., P.P.J and F.S. analysed the data. M.C. gave assistance in collecting tissue samples. C.C.L., P.P.J. and F.S. wrote the paper. P.P.J. and F.S. obtained funding. P.P.J and F.S. conceived the study and supervised it. All authors read and approved the submitted version.

## CONFLICT OF INTEREST

The authors declare no conflict of interest.

### Ethics Approval and Consent to Participate

Informed consent was obtained from enrolled patients and protocol was approved by Ministère de la Recherche (agreement n°: DC-2012-1598) and by local ethic committee (agreement n°: CB 2012/06).

