## Supplementary figures and images for "MCL-1 as a molecular switch between myofibroblastic and pro-angiogenic features of breast cancer-associated fibroblasts"

### Supp Figure 1

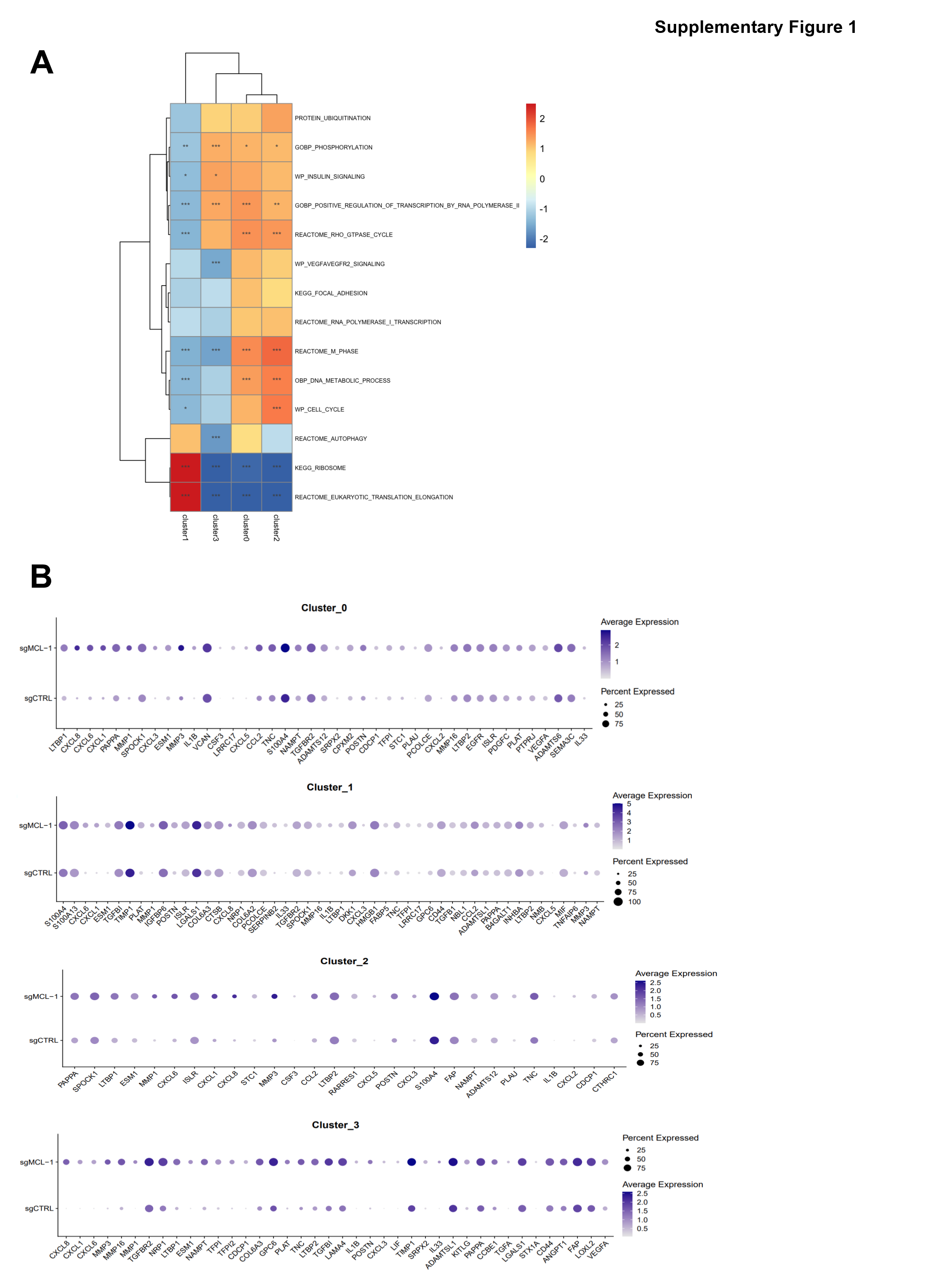

### Supp Figure 2

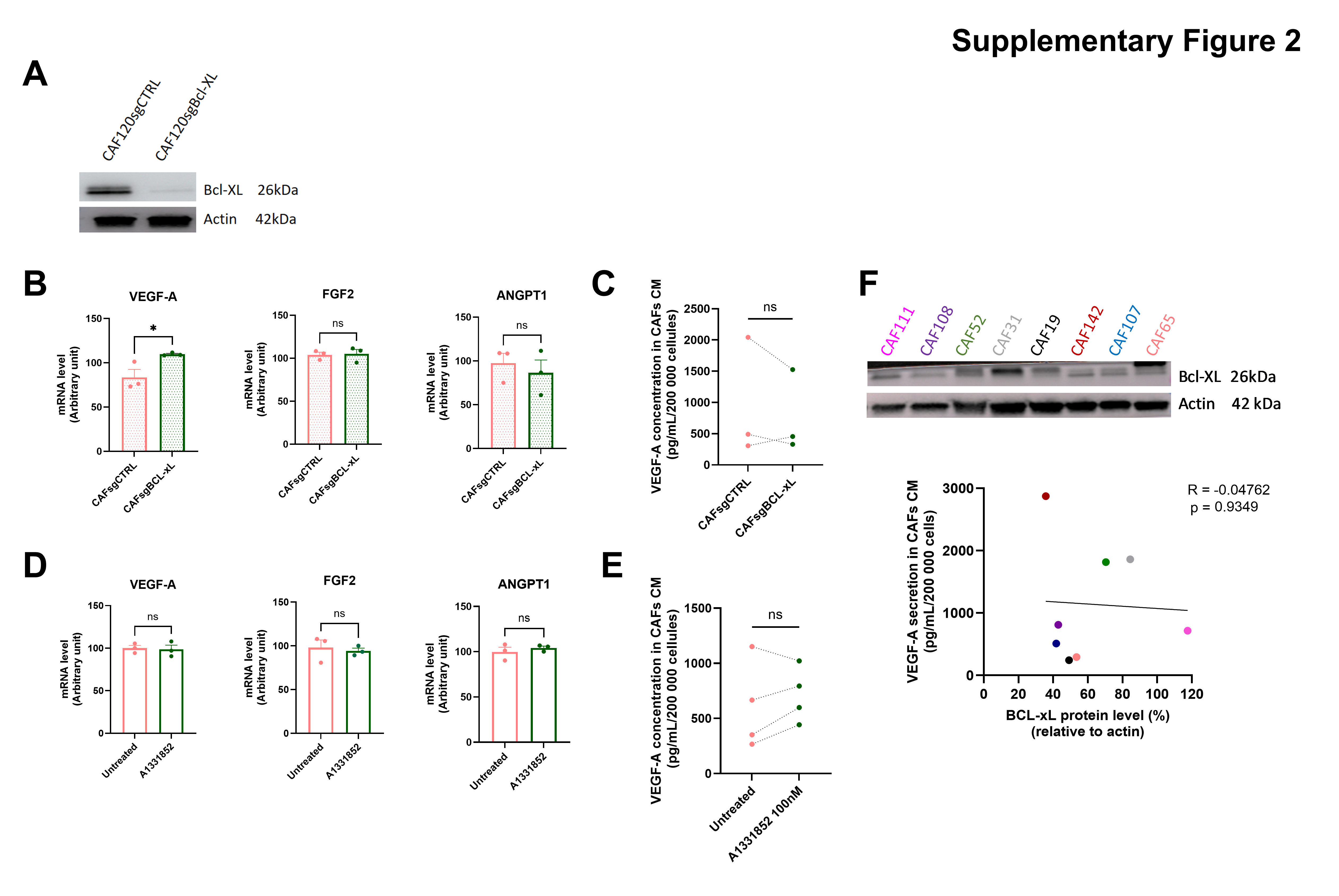

### Supp Figure 3

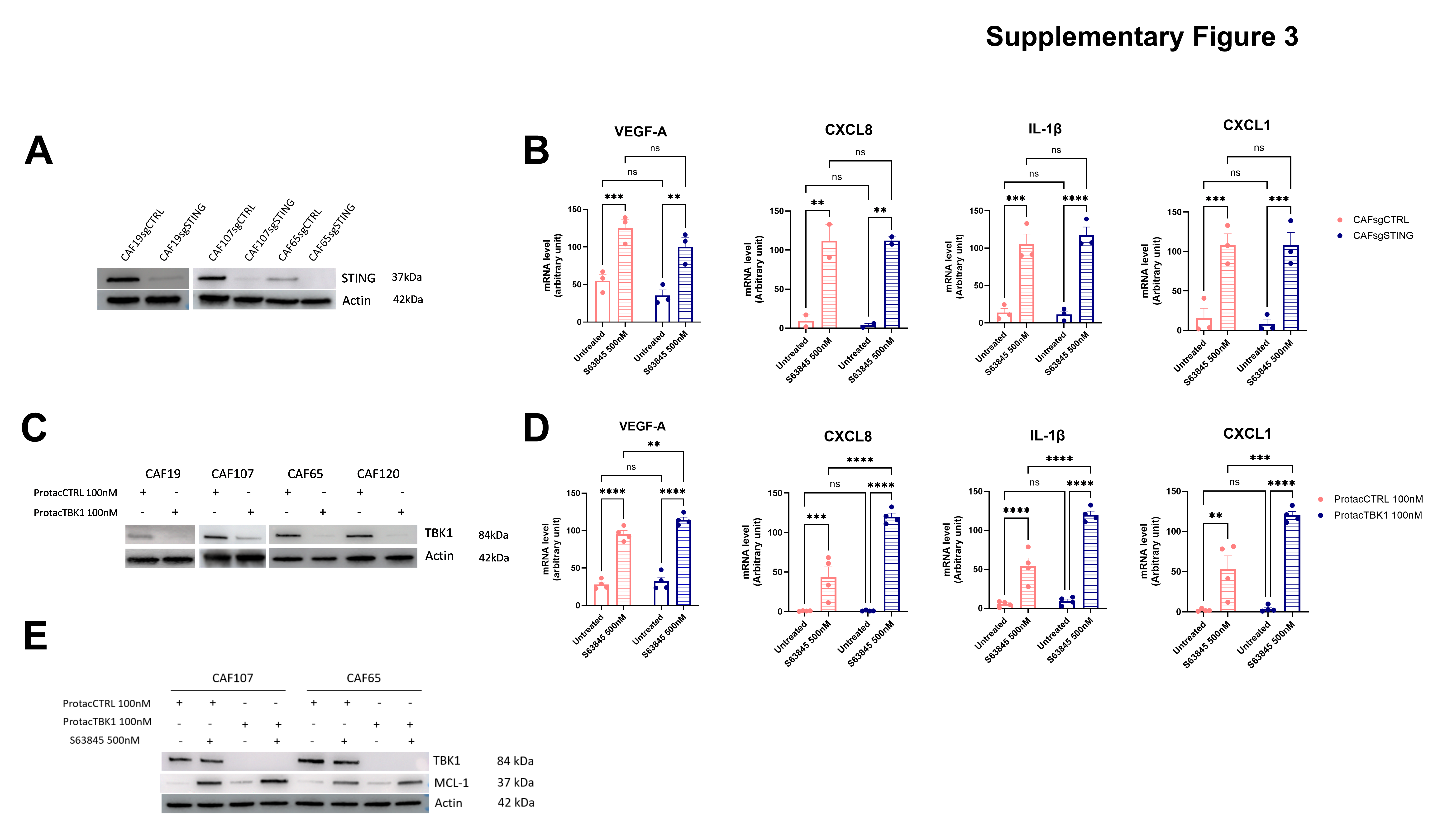

### Supp Figure 4

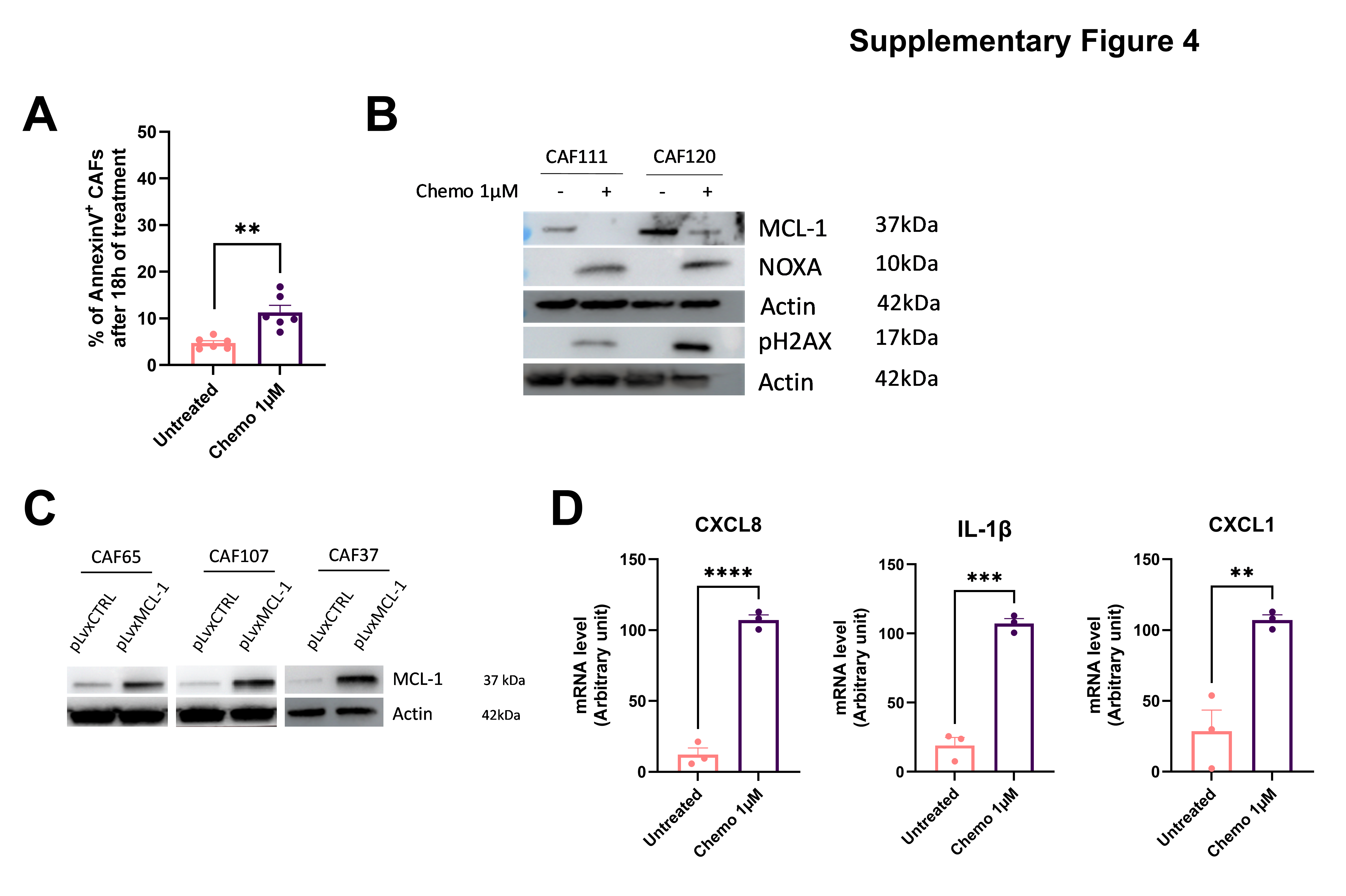
