## Supplementary material for "MCL-1 as a molecular switch between myofibroblastic and pro-angiogenic features of breast cancer-associated fibroblasts": Supp Table 1

|  |  |  | **Before QC** | | | **After QC** | | |
| --- | --- | --- | --- | --- | --- | --- | --- | --- |
|  | Total number of reads | Mapped Reads (%) | number of cells | Median UMI counts per cell | Median Genes per cell | number of cells | Median UMI counts per cell | Median Genes per cell |
| CAF51 KO CTRL | 86 752 918 | 96.5 | 2764 | 17916 | 4282 | 1778 | 24337 | 5029 |
| CAF51 KO MCL-1 | 45 988 891 | 97.3 | 3272 | 8778 | 2896 | 2205 | 10841 | 3328 |
| CAF65 KO CTRL | 174 692 440 | 96.9 | 5905 | 17481 | 4514 | 4920 | 18490 | 4664 |
| CAF65 KO MCL-1 | 227 249 947 | 97.4 | 8191 | 14774 | 4360 | 7177 | 15525 | 4486 |
| CAF107 KO CTRL | 138 693 334 | 97.4 | 5952 | 14288 | 4286 | 5074 | 14948 | 4419 |
| CAF107 KO MCL-1 | 224 266 763 | 97.6 | 8656 | 15752 | 4505 | 7802 | 16399 | 4618 |

**Supplementary Table 1: Quality Control Metrics for Single-Cell RNA Sequencing Data**
