## Supplementary material for "MCL-1 as a molecular switch between myofibroblastic and pro-angiogenic features of breast cancer-associated fibroblasts": Supp methods

**Supplementary METHODS AND MATERIALS**

**Apoptosis assay**

Cell death of bCAFs after 18h of chemotherapy was assessed using an Annexin-V FITC binding assay (Miltenyi #130-092-052) performed according to manufacturer’s instructions. Flow-cytometry analysis was performed on Accuri C6 Plus flow cytometer (BD Biosciences).

**Supplementary legends**

**Supplementary figure 1: A.** Enrichment analysis of gene sets preferentially expressed in cluster 0-3 in CAFs sgCTRL. Normalized enrichment scores and adjusted p-values were extracted per cluster, results were visualized as a heatmap displaying NES values, with significance annotations (* p ≤ 0.05, ** p ≤ 0.015, *** p ≤ 0.001). **B.** Dot plots of upregulated secreted factors (log2FC > 0.3, q-value < 0.01) in each cluster after MCL-1 gene silencing in bCAFs. The size of each dot represents the percentage of cells expressing the gene within the cluster, while the color intensity indicates the average expression level.

**Supplementary figure 2: Targeting of Bcl-XL doesn't modulate the pro-angiogenic factors secretion by bCAFs. A.** Bcl-XL protein expression level in bCAFs after gene silencing evaluated using western blots. Actin expression was used as loading control. **B.** qRT-PCR of VEGF-A, FGF2 and ANGPT1 mRNA in bCAF expressing BCL-Xl (sgCTRL) or not (sgBCL-Xl) (n=3). Student t-test, *P < 0.05, ns: not significant. **C.** VEGF-A quantification by ELISA in conditioned media (CM) from bCAFs after Bcl-XL gene silencing (bCAFsgBcl-XL) or not (bCAFsgCTRL). The bCAFs CM were generated during 72h in EGM2 (Endothelial Cell Growth Medium-2) medium supplemented with 1% of FBS (n=3). Student t-test, ns: non-significant. **D.** qRT-PCR of VEGF-A, FGF2 and ANGPT1 mRNA in bCAFs treated or not by A1331852 100 nM for 18 h (n=3). Student t-test, ns: not significant. **E.** VEGF-A quantification by ELISA in CM of bCAFs treated or not by A1331852 100 nM for 18 h Results were expressed as concentration (pg/ml) for 200 000 cells (n=4). Student t-test, ns: non-significant. **F.** Correlation (Spearman correlation coefficient r = −0.04762; p value = 0.9349) between VEGF-A secretion level in bCAFs CM and BCL-Xl protein level (relative to actin level) determined by western-blot analysis in 8 primary cultures of bCAFs.

**Supplementary figure 3:** **A.** STING protein expression level in bCAFs after gene silencing evaluated using western blots. Actin expression was used as loading control (n=3). **B.** qRT-PCR of VEGF-A, CXCL8, IL-1β and CXCL1 mRNA in bCAFs expressing STING (CAFsgCTRL) or not (CAFsgSTING) treated or not with S63845 for 18h (n=3). Two-way ANOVA, ****P < 0.0001, ***P < 0.001, **P < 0.01, ns: not significant. **C.** TBK1 protein expression level in bCAFs after 3h treatment with ProtacCTRL or ProtacTBK1 evaluated using western blots. Actin expression was used as loading control (n=4). **D.** qRT-PCR of VEGF-A, CXCL8, IL-1β and CXCL1 mRNA in bCAFs after treatment for 3h with ProtacCTRL or ProtacTBK1 before adding treatment for 18h with S63845 (n=4). Two-way ANOVA, ****P < 0.0001, ***P < 0.001, **P < 0.01, ns: not significant. **E.** TBK1 and MCL-1 protein expression levels in bCAFs after ProtacTBK1 or ProtacCTRL treatment for 3h before adding S63845 for 18h evaluated using western blots. Actin expression was used as loading control.

**Supplementary figure 4:** **A.** Apoptotic cell death of bCAFs after chemotherapy (1µM) for 18h in DMEM containing 1% FBS was measured by Annexin-V flow cytometry assay (n=6), Student t-test, **P<0.01. **B.** MCL-1, NOXA, pH2AX protein expression level in bCAFs after chemotherapy evaluated using western blots. Actin expression was used as loading control. **C.** MCL-1 protein expression level in bCAFs surexpressing MCL-1 (CAFpLvxMCL1) or not (CAFpLvxCTRL) were evaluated using western blots analysis. Actin expression was used as loading control. **D.** qRT-PCR of CXCL8, IL-1β and CXCL1 mRNA in bCAFs after treatment for 18h with chemotherapy (n=3). Student t-test, ****P < 0.0001, ***P < 0.001, **P < 0.01.
